# Reaching for domain-general syntax: Sentence processing and tool-use reach-to-grasp share neural patterns in the basal ganglia

**DOI:** 10.1101/2025.10.17.683107

**Authors:** Simon Thibault, Raphaël Py, Eric Koun, Roméo Salemme, Véronique Boulenger, Alice Catherine Roy, Claudio Brozzoli

## Abstract

Are actions organized like sentences? Recent evidence showed reciprocal transfer between tool use and syntactic comprehension, reflecting shared basal ganglia resources for action and language. The proposed mechanism is that embedding a tool into the motor plan increases the hierarchical structure of actions, paralleling the organization of sentences. If so, overlap with linguistic computations should emerge specifically during the initiation and/or reach-to-grasp phase, when tool embedding dynamically updates the relation with the target, rather than when the object is stably held in the tool grip. Forty French native speakers completed a sentence comprehension task involving object- and subject-relative clauses, and a motor task consisting of grasping and manipulating an object either with a tool or with the bare hand. Using both univariate and multivariate analyses, we found that the comprehension of syntactically complex sentences and the reach-to-grasp phase of tool use elicited overlapping and similar neural codes within the basal ganglia. This phase-specific convergence identifies the basal ganglia as a hub for domain-general hierarchical computations, unifying syntactic embedding for language and tool use.

## Introduction

Motor and communicative skills are exceptionally advanced in humans, as shown by our remarkable ability to use sophisticated tools and to convey meaning through complex linguistic structures[1–3]. Previous research has questioned whether these two refined skills co-evolved during human evolution and potentially rely on shared common cognitive resources[3–8]. The contribution of sensorimotor circuits to syntax, a crucial linguistic function, remains a matter of intense and perdurable debate[2,5,9–12]. Syntax is the function that governs the hierarchical organization of words into sentences, which mirrors the way actions can be assembled into hierarchically organized sequences[13,14]. The question therefore arises whether the organization of actions relies on computational processes akin to those underlying the hierarchical organization of sentences handled by syntax. Empirical evidence suggests this may indeed be the case. For instance, children with developmental language disorder exhibit disorganized kinematic patterns when performing everyday action sequences (i.e., reaching, grasping and lifting a bottle) compared to their neurotypical peers[15]. The functional relationship between action and syntax is also evident in neurotypical adults: observing daily life actions featuring structurally incorrect motor sequences triggers the same brain responses as those recorded when detecting syntactic violations in sentences[16].

In the same vein, dexterity in tool use correlates with the ability to produce complex syntactic utterances, even when controlling for cognitive load (i.e., working memory capacity)[17]. Furthermore, training participants to use a tool improves their comprehension of complex syntactic structures and vice versa[18,19]. Tool use indeed exemplifies the involvement of hierarchically organized sequences in the motor domain. To execute actions with a tool (e.g., grasping and manipulating a target object), one first needs to aptly grasp the tool with the hand, so to form a hand-with-tool unit (hereafter tooled hand). Ultimately, this process adds a hierarchical level in the action, where the tool becomes embedded within the manual motor plan and the tooled hand has to be handled towards the target object. The way tools are embedded within the overarching manual motor plan therefore bears similarities to linguistic structures, just as relative clauses can be embedded into main clauses (e.g., “*The writer [that the poet admires] reads the paper*”). Additionally, in both linguistic and motor domains, a unit can take a double function. In the object-relative sentence “*The writer that the poet admires reads the paper*”, the nominal phrase “*The writer”* serves as the subject of the main clause but also as the object of the center-embedded relative clause (“*the poet admires [the writer]*”). In the motor domain, the double function can similarly appear in tool use: the tool serves as the “subject” of the action, namely the effector that directly interacts with the target object; simultaneously it is the “object” of the hand. Taken together, these principles – embedding and double function – may rely on domain-general cognitive mechanisms for handling hierarchically organized structures, shared between language and action.

Recently, a fMRI study manipulating syntactic complexity in a sentence comprehension task, found that both tool use (i.e., inserting pegs into a pegboard with pliers) and syntactic processing engage subcortical areas, in particular the basal ganglia (BG)[18]. Such a finding is far from trivial as the BG are recognized as key nodes not only for sequencing actions[20,21], including those with tools[22–24], but also in syntactic processing for language[25–28]. This convergence therefore highlights the BG as the core neural hub for the domain-general cognitive mechanisms underlying tool use and syntax.

Despite the evidence that tool use and syntactic processes for language are co-localized within the BG[18], it remains nonetheless unclear which components of a tool-use action exhibit the strongest neural overlap with syntactic processing. Indeed, a movement can be finely decomposed into multiple phases, each reflecting distinct motor processes. Whether the different phases of tool-use actions engage shared neural resources with linguistic syntax to the same extent, or whether one phase of the action involves more common resources than the others, remains an open question. We here tackled this issue in a novel paradigm aiming to distinguish:

1. tool-use initiation, before any movement of the tooled hand.
2. the reach-to-grasp phase as the tooled hand approaches and establishes contact with the target object.
3. the object manipulation phase as the tooled hand moves the target object from one position to another.

These three phases entail different underlying processes. Initiation involves early motor planning with preparation of the different steps of the action sequence[29]. With a tool, this phase requires incorporating the tool in the body schema[30–32] and presumably tuning the sensory afferences related to this incorporation[33]. The reach-to-grasp phase then involves the first movement of the tooled hand, which may strengthen the process of tool incorporation and the reassignment of sensory afferences[34]. Reaching targets is known to be supported by online planning processes[35,36], requiring consideration of the target-object properties to adequately shape the tooled hand (or free hand) for grasping[15]. Lastly, the object-manipulation phase unfolds by implementing the motor plans set during the initiation and reach-to-grasp phases, yet still involving online control of the object to be displaced. Accordingly, the processes supporting tool embedding and its double function, which lead to a non-canonical action structure, are expected to be set and processed mainly during the initiation and/or reach-to-grasp phases. Indeed, these two phases require to dynamically update the relation between the tool and the target, as opposed to the manipulation where the target object is stably held in the tool’s grip. Therefore, we predict that the initiation and/or reach-to-grasp phases will recruit syntax-related processes to a greater extent than the object-manipulation phase (see the motor execution part in[18]). These processes will be reflected by tool-related brain activations closely relating to the activations observed for syntactic comprehension in language. In the current study, we thus tested competing hypotheses as to whether neural activity associated with complex syntactic processes shows more similarity to neural activity associated with (i) tool-use initiation, or (ii) reach-to-grasp, or (iii) both phases.

## Results

### Behavioral Performance in Syntactic and Motor Tasks

In a 3T MRI scanner, French native adult participants solved a task requiring them to process center-embedded relative clauses (Fig. 1A). Sentences relying on the same content words but featuring two different syntactic structures were visually presented: main clauses embedding a subject-relative clause (e.g., “The writer that admires the poet writes the paper”), or an object-relative clause (e.g., “The writer that the poet admires writes the paper”; Table S1A). After each sentence, participants had to judge if a test affirmation (e.g., “The poet admires the writer”; Table S1B) displayed on the screen was true or false with respect to the immediately preceding sentence. As expected, this task revealed longer response times for processing object-relative compared to subject-relative clauses [object-relative clauses = 1851 ± 74 ms vs. subject-relative clauses = 1578 ± 53 ms; χ^2^_(1)_ = 31.37; P < 0.001; Fig. 1B]. The sensitivity index (d′) was also smaller for object-relative clauses [1.57 ± 0.11 vs. subject-relative clauses = 2.27 ± 0.08; χ^2^_(1)_ = 33.63; P < 0.001; Fig. 1C], reflecting their increased processing complexity. All participants also performed a motor task in the scanner consisting of moving a peg (i.e., target object) from one side of a pegboard to the other with the bare hand (free hand hereafter) or with a tool (a 30-cm pair of pliers; Fig. 1D). The task was set-up to precisely measure timing information and break down the relevant phases of the motor behavior (i.e., initiation, reach-to-grasp and object-manipulation phases). To match task requirements between both tool-use and free-hand conditions, the participants were prompted to grab the tool placed on their chest before the start of a trial with a tool. In contrast, they were asked to return this tool to their chest before the start of a free-hand trial (see Methods, Video S1). In each condition, the initiation phase was defined as the time between the go signal and the moment the free or the tooled hand left the starting position (see Fig. S1). The reach-to-grasp phase was measured by the time elapsed between the onset of the movement and the grasping of the peg. Finally, the object-manipulation phase was indexed by the time spent pulling the peg out of its initial position and moving it to its final position on the pegboard. Participants were longer at initiating their movement with the tool rather than with their free hand [initiation times with the tool = 690 ± 15 ms, with the free hand: 656 ± 14 ms; χ^2^_(1)_ = 8.84; P = 0.003; Fig. 1E]. This is also true for the reach-to-grasp phase [reach-to-grasp times with the tool = 1499 ± 63 ms; with the free hand: 1104 ± 42 ms; χ^2^_(1)_ = 43.02; P < 0.001; Fig. 1F], and for moving and inserting the peg [object-manipulation times with the tool = 1648 ± 79 ms; with the free hand: 1423 ± 55 ms; χ^2^_(1)_ = 15.90; P < 0.001; Fig. 1G]. The participants completed seven runs in the MRI scanner for each of the two linguistic and motor tasks. As expected, behavioral analyses revealed that performance improved in both tasks along the different runs (Supplementary Text, Fig. S2).

**Figure 1:**
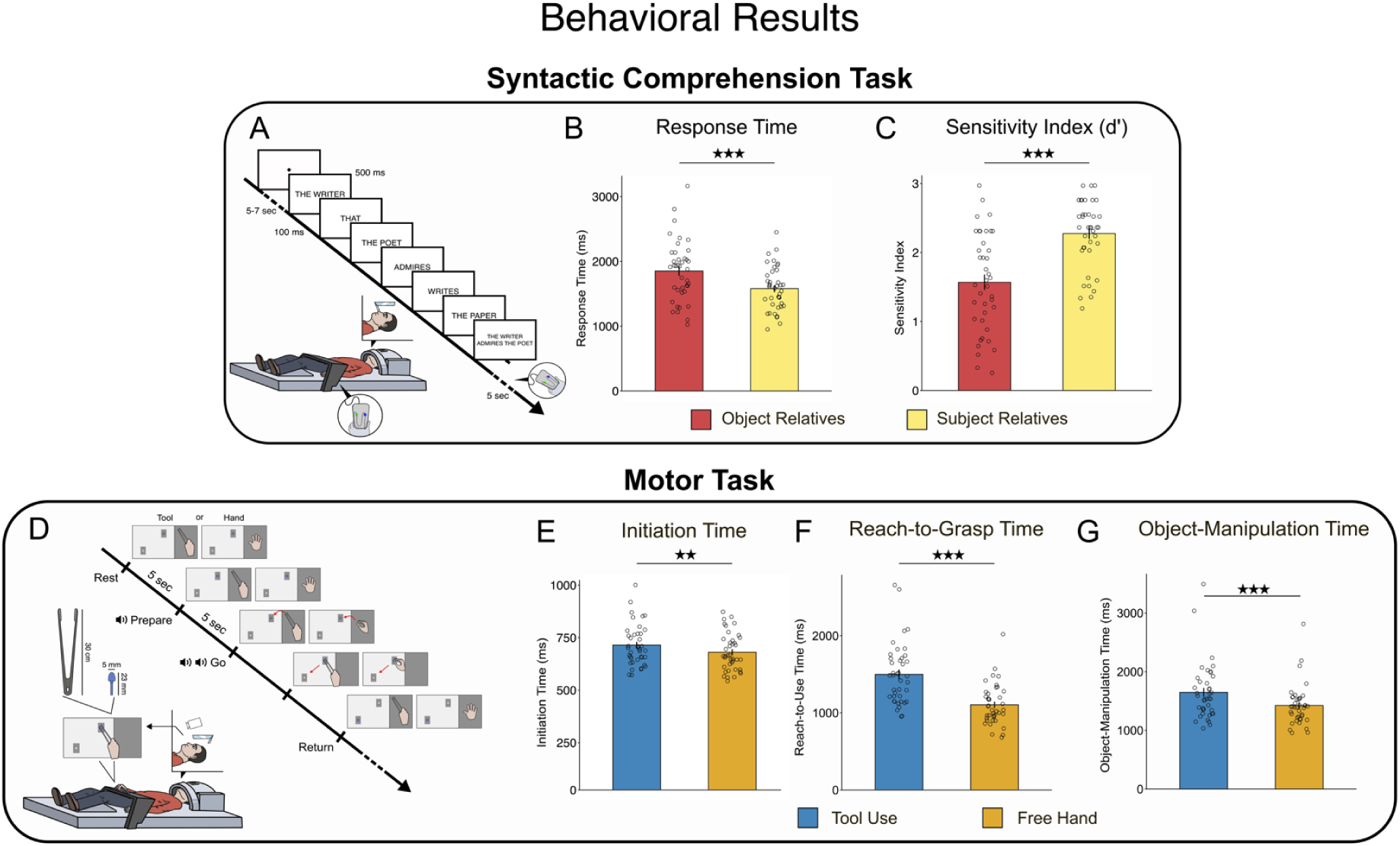
Syntactic comprehension task: experimental setup (A), response times (B) and sensitivity index (d′) (C). Behavioral performance is shown in red for object-relative clauses and in yellow for subject-relative clauses. Motor task: experimental setup (D), initiation time (E), reach-to-grasp time (F), object-manipulation time (G). Behavioral data for tool use is in blue and in orange for the free-hand condition. The stars represent the significance level (★★ p < 0.01; ★★★ p < 0.001). The error bars represent the standard error of the mean (SEM).

### Neural activations for syntactic processing and tool use

Both the syntactic and tool-use networks were assessed with a standard univariate whole brain approach. The effect of syntactic complexity was assessed by contrasting the neural activity elicited by processing object-relative compared to that elicited by subject-relative clauses during the sentence encoding phase (i.e., when the sentence was presented). Processing object-relative clauses significantly engaged both the left and right BG with peaks of activity localized in the caudate nuclei (cluster-corrected at p < 0.05; cluster forming threshold set at p < 0.001; Fig.2A; Table 1A). Notably, no significant difference in activity was found in the left inferior frontal gyrus (IFG) between object-relative and subject-relative clauses. Given the known role of the IFG in syntactic processing[37], we further investigated the activations within this region by conducting analyses in regions of interest (ROIs), namely the *pars opercularis* and the *pars triangularis* of the left IFG. In both ROIs, activations for object-relative clauses were significantly different from zero (t_s_ > 3.74; P_s_ < 0.001 uncorrected; P_s_ < 0.001 false discovery rate (fdr)-corrected; Cohen’s d_s_ > 0.59). This was also the case for the subject-relative clauses (t_s_ > 3.72; P_s_ < 0.001 uncorrected; P_s_ < 0.001 fdr-corrected; Cohen’s d_s_ > 0.59), consistent with the involvement of the left IFG in syntactic processing. In line with the whole-brain analysis, neither the *pars opercularis* nor the *pars triangularis* of the IFG showed sensitivity to syntactic complexity, as the activations for object-relative clauses were not significantly different from those elicited by subject-relative clauses in these ROIs (t_s_ < -0.07; P_s_ > 0.52 uncorrected; P_s_ > 0.53 fdr-corrected). We assessed the tool-use network by comparing the neural activity during tool use to that during free-hand movements, for each phase of the actions (initiation, reach-to-grasp and object manipulation). To get the most reliable estimate possible of neural activity during the initiation phase, we extracted the BOLD-activity from the 3-second period preceding the onset of the movement (i.e. time-locked to the participants’ reaction time). Initiating the action with the tool, compared to with the free hand, engaged a predominantly left-lateralized parieto-temporal network (cluster corrected at P < 0.05 like for the syntactic task; Fig. 2B; Table 1B). This network included a large cluster involving the supramarginal gyrus (SMG) and the superior parietal lobule (SPL), as well as a cluster in the lateral occipito-temporal cortex (LOTC). Additionally, a large cluster with a peak of activity in the right cerebellum was found, extending to the right inferior temporal lobe. Subcortical structures, including the left putamen, globus pallidus and thalamus, were also activated when initiating the movement with the tool.

**Figure 2.**
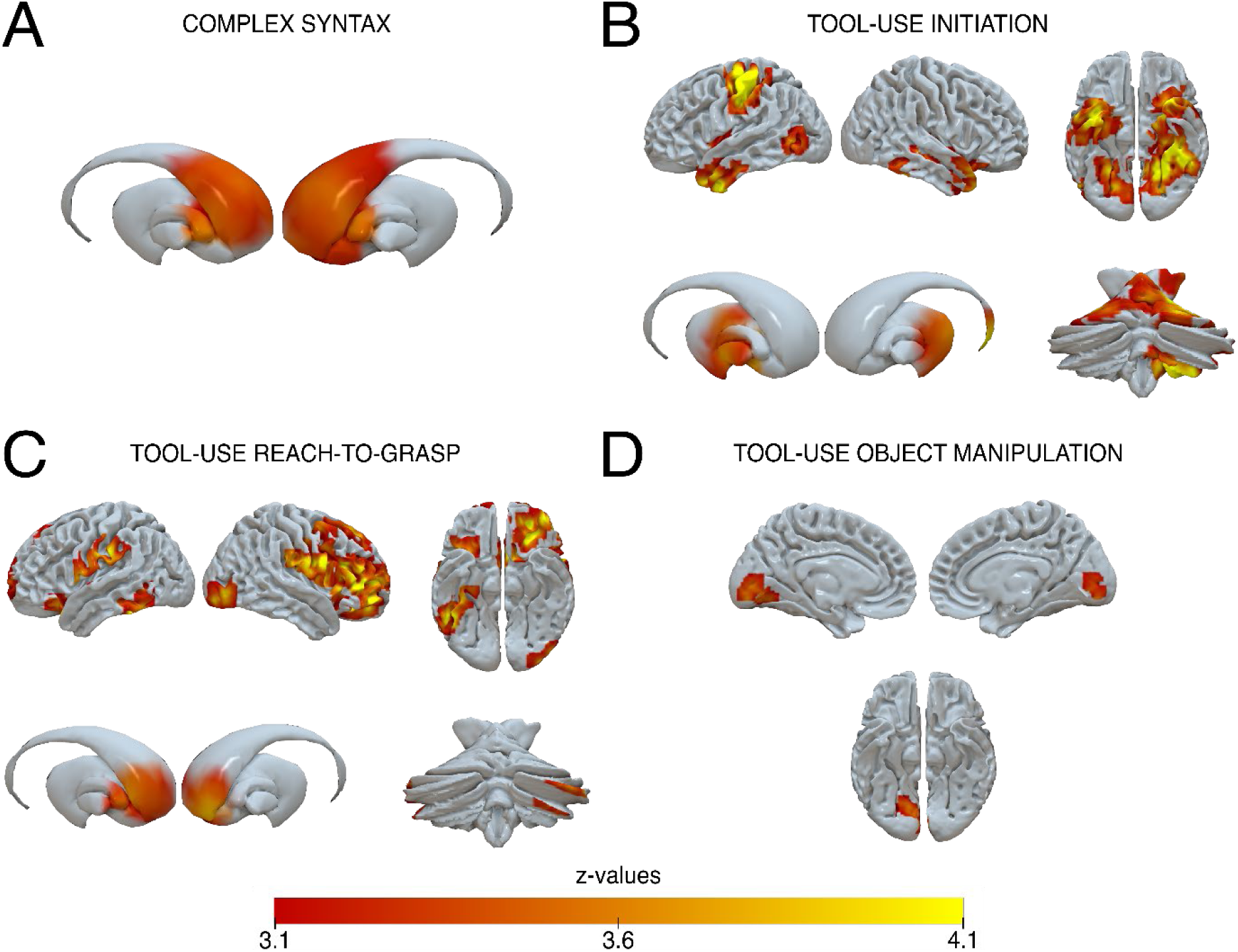
Neural activations for object-relative versus subject-relative clauses (A), tool-use initiation versus free-hand initiation (B), tool-use reach-to-grasp versus free-hand reach-to-grasp (C), tool-use object manipulation versus free-hand object manipulation (D). All maps were thresholded, clusters were formed at voxel-wise threshold of P = 0.001 and corrected at the cluster level at P = 0.05. The localization of the activations is described in Table 1.

**Table 1.**
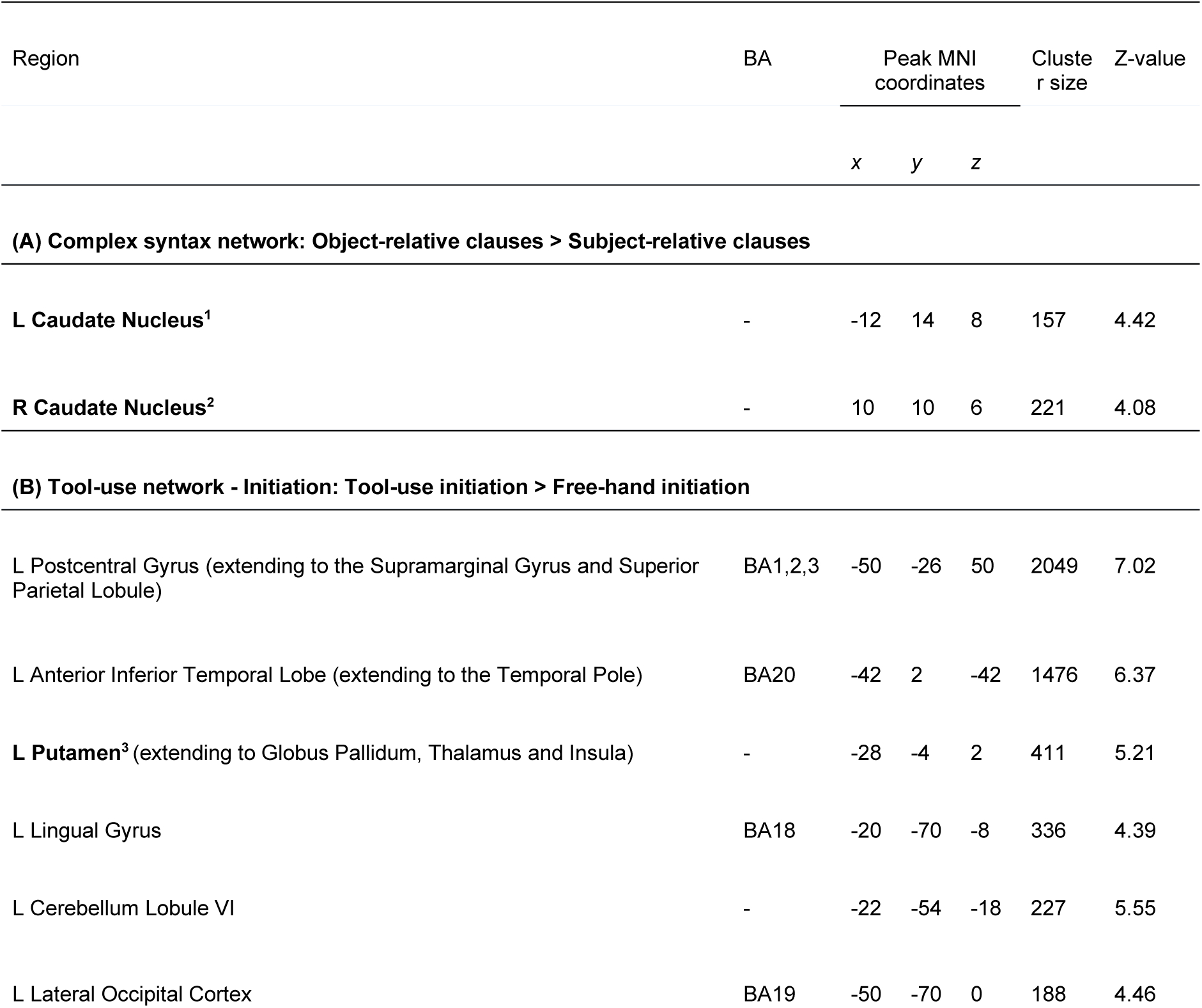
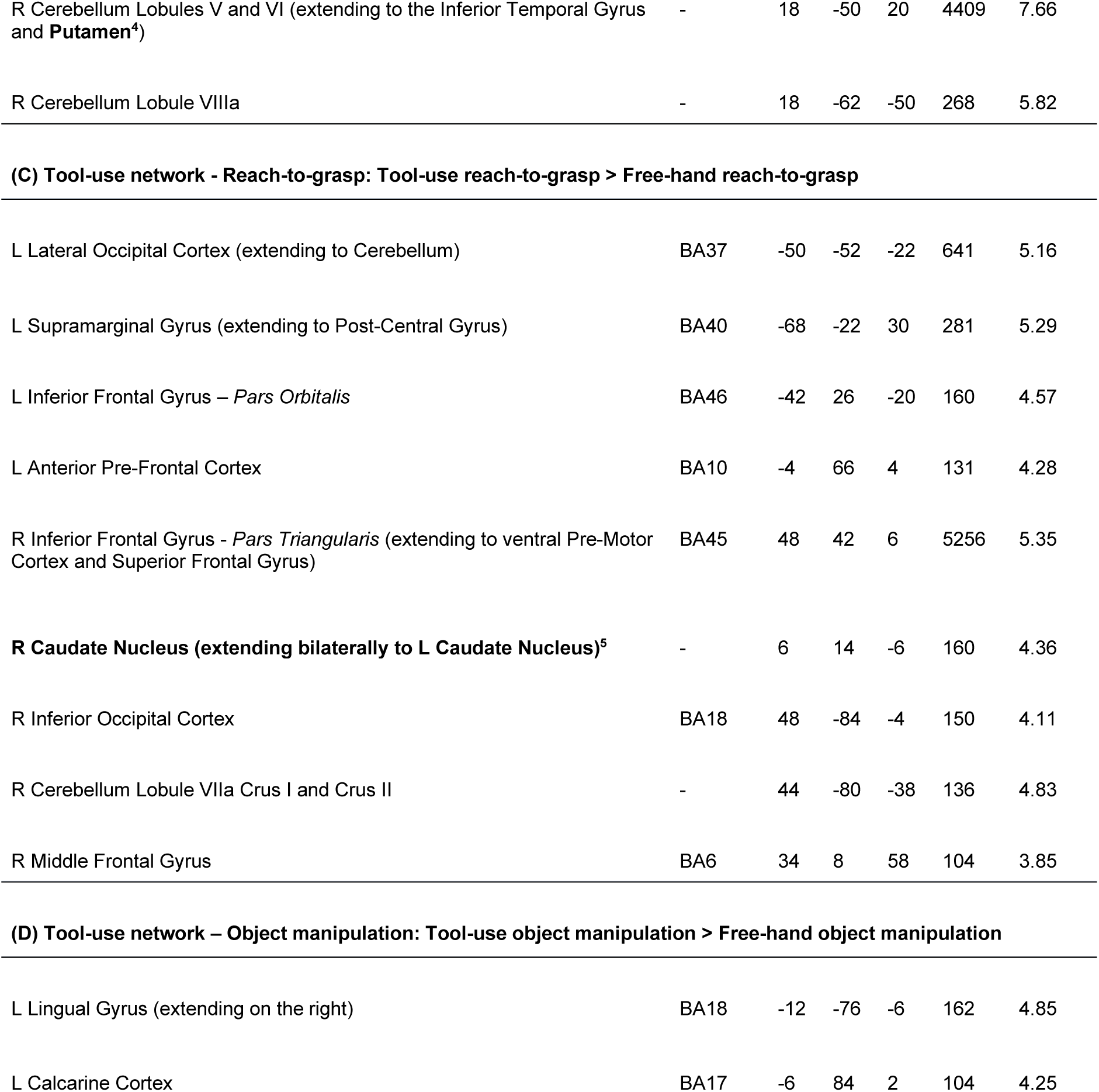
Brain regions activated for A) object-relative versus subject-relative clauses, B) tool use versus free hand in the initiation phase, C), tool use versus free hand in the reach-to-grasp phase D) tool use versus free hand in the object-manipulation phase. From left to right are reported the names of the regions in which the clusters were identified, the Brodmann Area (BA), the Montreal Neurological Institute (MNI) coordinates for the peak of activity, the cluster size in number of voxels and the statistics with the Z-values. The clusters highlighted in bold were used for regions-of-interest (ROIs) analyses.

The reach-to-grasp phase, namely from movement onset to peg grasping, with the tool compared to the free hand involved a bilateral network including left fronto-parietal regions (cluster corrected at P < 0.05; Fig. 2C; Table 1C), with activity in the left SMG and IFG (*pars orbitalis*). In the right hemisphere, a large cluster also displayed activity in the IFG (*pars opercularis*). Additionally, subcortical activity related to reaching and grasping with the tool (compared to the free hand) was observed bilaterally in the caudate nucleus. Finally, using the tool for manipulating the peg (i.e. pulling out, transporting and inserting the peg to a different location) induced distinct neural activity in the left inferior occipital gyrus (IOG, cluster corrected at P < 0.05; Fig. 2D; Table 1D) relative to the free-hand object manipulation.

### Overlap between syntax and tool-use neural networks

We further examined the co-localization of brain activity between tool use and syntax in the BG, regions for which we had a priori hypotheses. These ROIs were selected from the univariate whole-brain analyses presented in the previous section. For each region activated by complex syntactic processing (i.e., object-relative vs. subject-relative clauses), we tested whether tool use elicited greater neural activity (i.e. higher β estimates) than free-hand actions for each phase separately. Conversely, for clusters activated during each tool-use phase (i.e., tool use vs. free hand), we tested whether object-relative clauses elicited greater neural activity than subject-relative clauses. In total, we selected two clusters significantly associated with complex syntactic processing (i.e. object-relative clauses), both located in the left and right caudate nuclei (Table 1A^1,2^). For tool use, we identified three significant clusters: two clusters in the left and right putamen activated during tool-use initiation (Table 1B^3,4^), and one spanning bilaterally over the caudate nuclei (i.e., contiguous voxels in the left and right caudate) activated during the reach-to-grasp phase of tool use (Table 1C^5^). No cluster was selected from the object-manipulation phase, as this phase did not reveal stronger activations for tool-use than free-hand actions in the BG.

Considering the clusters activated by complex syntactic processing, we found greater activity for the reach-to-grasp phase with the tool compared to the free hand in the bilateral caudate nuclei [in the left: tool-use reach-to-grasp β = 1.93 ± 1.39; free-hand reach-to-grasp β = -2.55 ± 1.64; t_(39)_ = 2.51; P = 0.008 uncorrected; P = 0.02 fdr-corrected; Cohen’s d = 0.40; Fig. 3A; in the right: tool-use reach-to-grasp β = 3.58 ± 0.77; free-hand reach-to-grasp β = -0.30 ± 1.73; t_(39)_ = 1.91; P = 0.03 uncorrected; P = 0.03 fdr-corrected; Cohen’s d = 0.30; Fig. 3B]. In contrast, no significant differences were found in these subcortical regions when comparing tool-use and free-hand actions during the initiation and the object-manipulation phases [t_s_ < -1.79; P_s_ > 0.95 uncorrected; P_s_ > 0.99 fdr-corrected, Fig. S3].

**Figure 3:**
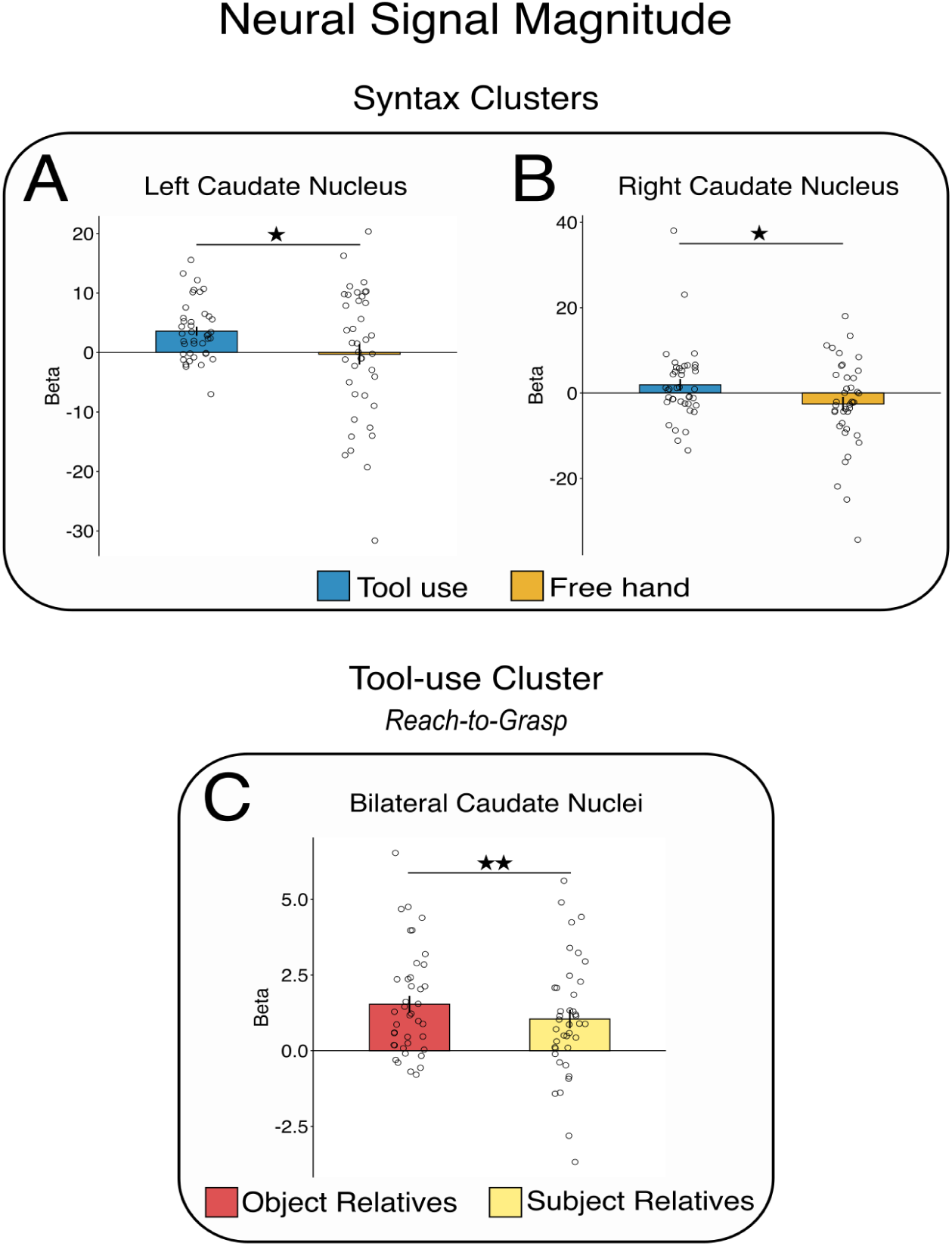
Motor-averaged β in complex syntax ROIs, namely the left (A) and right (B) caudate nuclei clusters activated by object-relative vs. subject-relative clauses. In these clusters, neural activity for tool-use reach-to-grasp (blue) was on average significantly stronger than free-hand reach-to-grasp activity (orange). Syntax-averaged β in the tool-use ROI, namely the caudate cluster activated bilaterally for tool-use reach-to-grasp is shown in (C). In this cluster, neural activity for object-relative clauses (red) was on average significantly stronger than the one elicited by subject-relative clauses (yellow). Each black circle represents a unique participant. The stars represent significance level after corrections for multiple comparisons (★ p < 0.05; ★★ p < 0.01). The error bars represent the standard error of the mean (SEM).

When considering the clusters significantly activated by tool use, we found stronger activity for object-relative compared to subject-relative clauses in the bilateral caudate nuclei involved in the reach-to-grasp phase [object-relative clauses = 1.54 ± 0.27; subject-relative clauses = 1.05 ± 0.30; t_(39)_ = 2.37; P = 0.01 uncorrected; P = 0.03 fdr-corrected; Cohen’s d = 0.37; Fig. 3C]. However, we did not observe any difference between object-relative and subject-relative clauses, neither in the left nor the right putamen involved in the tool-use initiation phase [t_s_ < -0.15; P_s_ > 0.56 uncorrected; P_s_ > 0.84 fdr-corrected; Fig. S3]. These ROI-based findings were corroborated by a whole-brain conjunction analysis revealing a cluster of shared voxels in the left caudate when processing object-relative clauses and reaching and grasping with the tool (see Fig. S4, Table S2).

### Tool use and complex syntax activations display similarities in their neural representations

To thoroughly test whether tool use and complex syntactic processing share neural resources within the BG, we assessed the similarity of their neural representations. We ran multivariate analyses to compare the fine-grained spatial organization of the brain activations elicited by the syntactic and the motor tasks. Significantly similar patterns of activation between tool use and complex syntactic processing would indicate the existence of common neurofunctional resources. We conducted this analysis within the same ROIs activated by tool use or complex syntax, as identified in the previous univariate analyses based on signal magnitude: 1) the left caudate and 2) the right caudate activated by complex syntactic processing; 3) the left putamen and 4) the right putamen activated by tool-use initiation; and 5) the bilateral caudate activated by reach-to-grasp with the tool. To assess the extent of neural patterns similarity, we computed the cosine similarity between syntactic and motor conditions, within these five ROIs. A higher cosine similarity indicates greater similarity between neural patterns. We measured (i) the cosine similarity between the neural patterns elicited by object-relative clauses and tool-use. As controls, we also computed the similarity between (ii) subject-relative clauses and tool-use and (iii) object-relative clauses and free-hand.

In the syntax-activated clusters, we found significantly stronger cosine similarity between patterns elicited by object-relative clauses and tool-use reach-to-grasp in the right caudate nucleus [cos = 0.13 ± 0.02; Fig. 4B in purple], compared to both the similarity between subject-relative clauses and tool-use reach-to-grasp [cos = 0.10 ± 0.02 ; t_(39)_ = 2.95; P = 0.003 uncorrected; P = 0.008 fdr-corrected; Cohen’s d = 0.47; Fig. 4B in green], and between object-relative clauses and free-hand reach-to-grasp [cos = 0.06 ± 0.03 ; t_(39)_ = 2.10; P = 0.02 uncorrected; P = 0.04 fdr-corrected; Cohen’s d = 0.33; Fig. 4B in orange].

**Figure 4:**
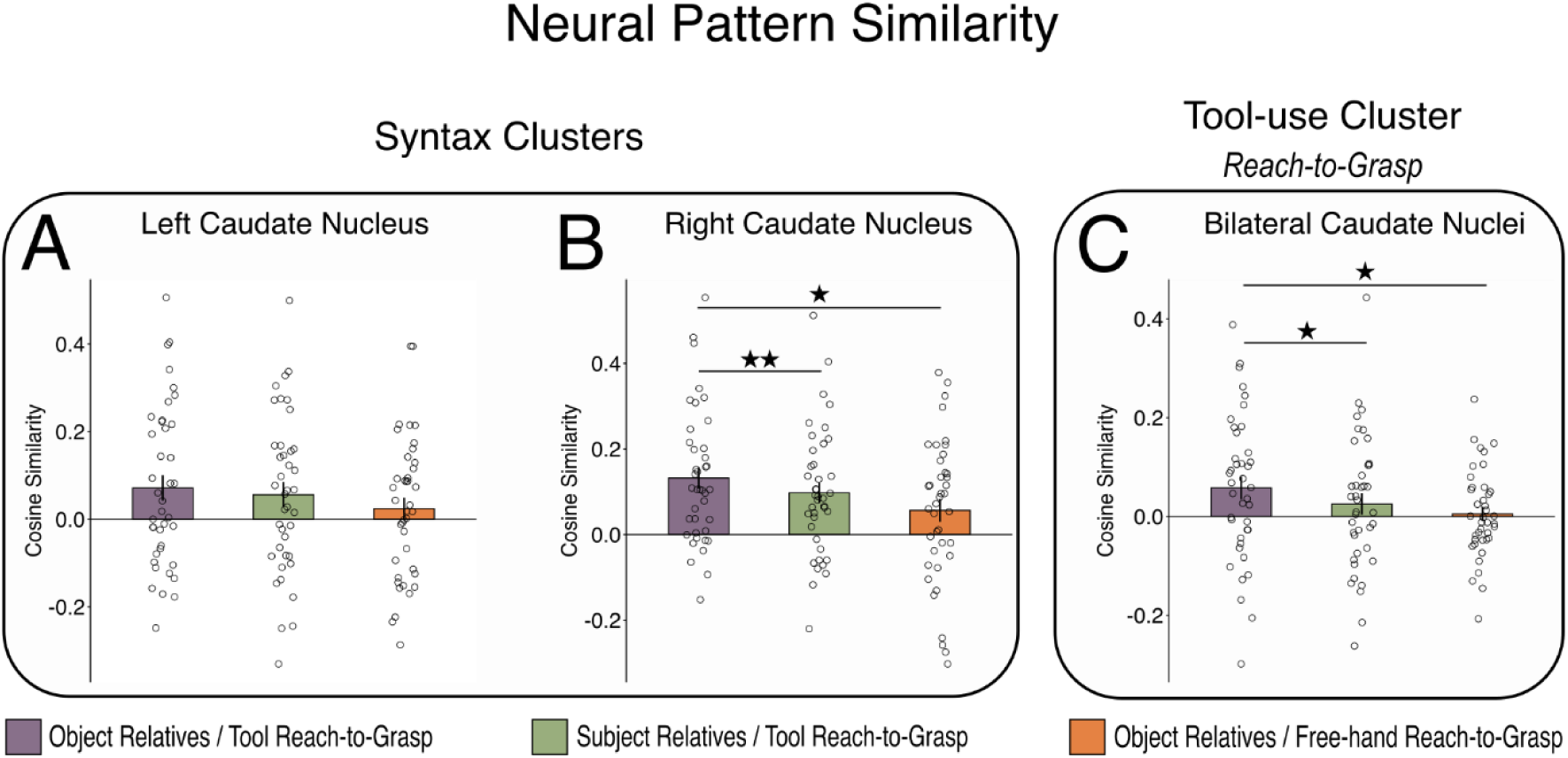
Cosine similarity in the left (A) and right (B) caudate nuclei clusters activated for object-relative clause processing and in the bilateral caudate cluster (C) activated by tool-use reach-to-grasp. The higher the cosine similarity, the more similar the neural patterns of the conditions in the pair. The cosine similarity between object-relative clauses and tool-use reach-to-grasp patterns is shown in purple. The cosine similarity between subject-relative clauses and tool-use reach-to-grasp patterns is represented in green, and the cosine similarity between object-relative clauses and free-hand reach-to-grasp patterns is in orange. Each black circle represents a unique participant. The stars represent significance level after corrections for multiple comparisons (★ p < 0.05; ★★ p < 0.01). The error bars represent the standard error of the mean (SEM).

In contrast, in the left caudate nucleus also activated by complex syntax, the similarity between patterns elicited by object-relative clauses and tool-use reach-to-grasp [cos = 0.07 ± 0.03; Fig. 4A in purple] was not significantly different from the similarity either between subject-relative clauses and tool-use reach-to-grasp [cos = 0.06 ± 0.03 ; t_(39)_ = 1.09; P = 0.14 uncorrected; P = 0.14 fdr-corrected; Fig. 4A in green], or between object-relative clauses and free-hand reach-to-grasp patterns [cos = 0.02 ± 0.03 ; t_(39)_ = 1.37; P = 0.09 uncorrected; P = 0.09 fdr-corrected; Fig. 4A in brown].

In these two syntactic clusters, we also analyzed the pattern similarity between the syntactic activations and the motor activations for the initiation and object-manipulation phases, respectively. The pattern similarity was not significantly stronger for object-relative clauses and tool-use initiation compared to either subject-relative clauses and tool-use initiation, or object-relative clauses and free-hand initiation [t_s_ < -0.16; P_s_ > 0.95 uncorrected; P_s_ > 0.99 fdr-corrected, Fig. S5]. The same was found when considering the activation patterns for object manipulation and their similarity with the syntactic activation patterns [t_s_ < - 0.16; P_s_ > 0.67 uncorrected; P_s_ > 0.67 fdr-corrected, Fig. S6].

In tool-use activated clusters, for the bilateral caudate nuclei involved in tool-use reach-to-grasp, we found that the neural patterns elicited by object-relative clauses and by tool-use reach-to-grasp [cos = 0.06 ± 0.02; Fig. 4C in orange] were significantly more similar compared to those elicited by subject-relative clauses and tool-use reach-to-grasp [cos = 0.03 ± 0.02; t_(39)_ = 2.01; P = 0.03 uncorrected; P = 0.04 fdr-corrected; Cohen’s d = 0.32; Fig. 4C in green], and to those elicited by object-relative clauses and free-hand reach-to-grasp [cos = 0.01 ± 0.01; t_(39)_ = 2.02; P = 0.03 uncorrected; P = 0.04 fdr-corrected; Cohen’s d = 0.32; Fig. 4C in orange]. In contrast, in the left and right putamen, which were involved in tool-use initiation, we did not find stronger pattern similarity between object-relative clauses and tool-use initiation compared to either subject-relative clauses and tool-use initiation, or object-relative clauses and free-hand initiation [t_s_ < 0.62; P_s_ > 0.26 uncorrected; P_s_ > 0.99 fdr-corrected, Fig. S5].

## Discussion

Neural co-localization of syntax and tool use in the basal ganglia (BG) has pointed to shared neurocognitive resources[18]. In the current study, we systematically dissected the distinct phases of tool-use action – namely initiation, reach-to-grasp and object-manipulation – uncovering how these processes map onto neural activity and show similarities with syntactic processing. Our key finding is that the neural co-localization in the BG between tool use and complex syntactic processing is observed specifically for the reach-to-grasp phase, but not for the initiation nor the object-manipulation phases. Moreover, within part of these shared clusters, we show a similar fine-grained spatial distribution of the neural patterns, providing evidence for tight functional links between complex syntax and the reach-to-grasp phase of tool use. These findings indicate that the motor phase of the action where the tool, held in the hand, needs to be articulated with the target object engages neural activity akin to that involved in complex syntactic comprehension. Processing hierarchically organized structures in both action and language therefore relies on common neural bases, pointing to a core, domain-general syntactic function.

In the language domain, previous studies have shown that neurological conditions affecting the basal ganglia are associated with comprehension deficits for complex syntactic structures[38,39]. These findings are further corroborated by neuroimaging studies showing activation of the basal ganglia during the processing of syntactically complex sentences[25,27,40,41]. In the motor domain, it is well established that the basal ganglia play a critical role for action sequences[20,21,42,43]. Importantly, these subcortical structures are also activated for tool-related actions[18,22–24]. The involvement of the basal ganglia might stem from the more complex hierarchical structure introduced by tool-mediated actions compared to manual actions. Indeed, using a tool requires integrating one further level into the action structure when compared to free-hand action. This is reflected by the need to translate the tool (primarily an object) into an effector to interact with a secondary object, the target. As a result, the overall process reflects the syntactic organization of interdependent elements. The analogy between syntactic processes and tool use can indeed be extended to the structure of sentences and tool-related actions. Compared to subject-relative clauses (e.g., “the writer that admires the poet reads the paper”) where words follow a linear, canonical order (i.e., subject – verb – object for French and English languages), object-relative clauses (e.g., “The writer that the poet admires reads the paper) feature a non-canonical structure in which this linear order is disrupted, with the object preceding the verb, and the subject following it. A similar distinction may be drawn in the motor domain: grasping an object involves a direct relationship between the hand and the target object, reflecting a canonical structure. By contrast, tool use implies an indirect, hierarchical (non-canonical) relationship, whereby the hand acts indirectly on the target object by the means of a tool. Processing such non-canonical structures may rely on domain-general cognitive mechanisms shared between language and action.

An alternative explanation may be that the shared activations patterns observed in the BG result from a higher cognitive load associated with tool-use and the syntactic processing of object-relative clauses. However, previous investigations argue against this interpretation. Carefully controlled sentence-processing paradigms have indeed shown that syntactic structure comprehension can be dissociated from executive demands, including working memory[44,45]. With respect to our specific protocol, a previous neuroimaging study compared the neural activations elicited by tool use and object-relative sentences and further contrasted them with activations elicited by varying levels of difficulty in a working memory task (one-vs. three-back task)[18]. The results revealed a co-localization of tool-use related and syntactic processing activations, but no overlap with working memory activity. This pattern ruled out the possibility that the observed overlap between tool use and syntactic processes reflected shared neural mechanisms driven by working memory or higher general cognitive load. Moreover, a recent work[19] reported improved syntactic comprehension after tool-use training, irrespective of the complexity of the motor task (i.e., moving three target objects vs. one), suggesting that additional cognitive resources do not modulate this transfer effect. Similarly, a study examining the relationship between tool use and syntactic processing in adults found that tool-use dexterity predicts syntactic production abilities, but not verbal working memory performance [17]. Altogether, these findings suggest that cognitive control is unlikely to constitute a core resource for processing hierarchical structures.

Our results findings indicate that more than the other phases, the reach-to-grasp phase of tool use may rely on syntactic operations required to embed the tool in the overarching motor plan and integrate its double function in the action (i.e., tool as both a “*subject*”/effector of the action and an “*object*” of the hand). During this phase, the manual sensorimotor program must be updated on-line to functionally account for the presence of the tool in the hand and prepare the interaction with the target object. As a result, this may exacerbate the need to properly encode the hierarchy among interdependent elements of the sensorimotor sequence, thereby enabling the coordinated interaction between the hand, the tool and the target object. Our findings also suggest that, during the initiation phase, the on-line encoding of the relationships between these elements of the action may be less critical.

Combining different objects together to form more complex ensembles is a non-trivial aspect of cognition that has been studied with the nesting-cup paradigm. This paradigm has been classically used to study the abilities to generate hierarchically organized action structures in different species and at different life periods[46]. In its simplest version, it requires assembling three cups of different sizes together (i.e., the smallest ones into the largest cup). Two main strategies are usually observed to solve the task: the “*pot*” and the “*subassembly*” strategies. In the “*pot*” strategy, the medium cup is first inserted into the largest cup before the small cup gets inserted into the medium one. In contrast, the “*subassembly*” strategy consists in first combining the smallest cup with the medium one together, before combining this subset of cups with the largest cup. Human adults usually rely more on the “*subassembly*” strategy than chimpanzees[47]; infants develop this strategy only after 18 months of life[46,47]. The (more complex) “*subassembly*” strategy is considered to involve hierarchically organized combinatorial activity, as opposed to the (simpler) “*pot*” strategy consisting of serially adding each cup to the largest one, starting from the biggest[48]. Accordingly, the subassembly strategy has been considered as a precursor for tool use[47] and potentially syntax in language[49].

Hierarchical processing of different elements within a sensorimotor sequence[50] is emphasized during the reach-to-grasp phase of action with objects. A previous study revealed that neurotypical children required to grasp and lift a bottle, were able to adapt kinematic parameters in the preceding reach-to-grasp phase when the weight of the bottle was known beforehand, so to ease the subsequent lifting phase[15]. Interestingly, this functional anticipatory behavior was not observed during the reach-to-grasp phase in children with a Developmental Language Disorder (DLD), who show impaired syntactic abilities. This finding highlights the crucial importance of the reach-to-grasp phase for integrating the interdependence between each element of the action. In line with our neuroanatomical findings, previous reports suggested that children with a DLD show dysfunctional basal ganglia activity[51,52]. Thus, the ability to process interdependence between elements in action may require the same cognitive function as that engaged in processing the interdependence for syntax in language. This idea found support in Patricia Greenfield’s observations of infants’ behavior[53]. She suggested that language acquisition and advanced motor skills rely on domain-general processes enabling elementary combinations of units (phonemes/words for language, and objects for action). These elementary combinations can serve as a scaffold for building more complex structures, such as those observed for tool use in action and for embedded sentences in language[53].

The current findings specifically highlight the role of the basal ganglia in processing complex structures in both the language and action domains. It has been proposed that the BG (together with the frontal cortex) are part of a procedural system supporting actions as well as the learning of regularities in language[52,54–56]. Understanding complex structures in language, such as center-embedded object relative clauses, requires managing the hierarchical relations between sentence components. Similarly, tool use involves managing the relationship between each component of the action sequence. Processing such complex, structured regularities in language and action thus likely relies on domain-general processes that are shared across both domains. These domain-general processes may be supported by a procedural system hosted in the basal ganglia, handling domain-general syntactic-like computations[52,54–56]. The involvement of the basal ganglia in such domain-general processes may reflect their ancient phylogenetic character[57,58]. In numerous species, these subcortical structures are indeed involved in behaviors requiring the combination of simple elements into longer, structurally organized chunks[42]. The BG may therefore have been one region that has played a critical role in the emergence of language and tool use in humans. Importantly, as part of different cortico-subcortical loops[59,60], their precise contribution needs to be further investigated, to understand how they support shared or domain-specific networks for language and tool use.

As suggested, the left inferior frontal gyrus (IFG) could also fulfill a domain-general function in both language and action[61]. Neuroimaging studies reported stronger activity in the left IFG when processing complex structures in both the language[37] and action domains[13]. In neurotypical adults, reading sentences and using tools indeed both activate the left IFG[4]. People with agrammatic aphasia following a lesion in Broca’s area are furthermore impaired in ordering biological action events[62]. In the current study, however, we found comparable left IFG activity (i.e., *pars opercularis* and *pars triangularis*) for more (object relatives) and less complex (subject relatives) syntactic structures. The lack of difference between the two syntactic conditions in this cortical region might stem from the fact that their general structure remained comparable (center-embedding as opposed to coordinated structures) and with very little variation in their lexical content[63]. Given the lack of stronger IFG-related activations for object-relative clauses compared to subject-relative clauses, as well as for tool use compared to free-hand actions, we suspect this region does not really act as a supramodal processor for hierarchically organized structures. This observation is in line with previous work showing that syntax in language and action engage distinct portions of the inferior frontal area[18,64,65]. Activations for syntax in language usually involve more anterior regions such as the *pars opercularis*, while more posterior activations within the ventral premotor cortex are observed for manual actions and tool use.

The current findings of neural co-localization between syntax in language and tool use should also be considered with respect to recent work suggesting that structures in language and action are different (e.g.[66,67]). These studies argue for a high degree of specialization in brain areas dedicated to language, suggesting that the sophistication of linguistic structures does not extend to action structures[66]. As a result, such complex linguistic structures would be subserved by domain-specific brain areas such as the IFG which would have expanded anteriorly under the pressure to handle linguistic processes[67]. However, substantial evidence indicates that the IFG is also involved in a wide range of cognitive processes[68,69], casting doubt on its domain-specificity for language, which remains an open question. We propose that language recruits more elementary processes to build highly specific structures. These fundamental processes are shared with action, enabling the combination of elements into more complex structures. Our findings suggest that phylogenetically older structures, such as the basal ganglia, may contribute to domain-general computations for processing hierarchical structures. Thus, syntax may rely on elementary computations subserved by the basal ganglia[52,54–56], rather than solely on the expansion of neural resources within the IFG[67].

## Materials and Methods

### Participants

A sample of 46 adult participants took part in this study. Three of them did not fulfill the pre-test requirements (see Procedure) to be enrolled in the fMRI session and were excluded. Three additional participants dropped out after the pre-test session and did not show up for the fMRI session. Overall, 40 participants underwent the fMRI session and were considered for the subsequent analyses. This sample of 40 participants (females = 20) had the following demographic characteristics: age (mean ± SD) = 22.98 ± 3.25 years; education years after high school = 2.98 ± 1.31 years; handedness = 0.94 ± 0.08[70]. All the participants gave their consent to take part in the study. The procedure complied with the Declaration of Helsinki and has been validated by a national ethics reviewing committee (CPP OUEST IV).

### Syntactic Task

A two-alternative forced choice task (2-AFC) assessed the syntactic abilities supporting the comprehension of sentences with the same content words but featuring different structures: main clauses with center-embedded subject relative and center-embedded object relative clauses. Table S1A offers examples for each condition, and the entire material is available at https://osf.io/6adfn. The content words included in the sentences were controlled for word frequency and number of syllables from the Lexique 3.80 database[71], as well as for the gender (feminine or masculine nouns) of the subjects and objects of the described action. Each sentence was presented using rapid serial visual presentation (RSVP) in six consecutive segments displayed in the middle of the screen for 500 ms, interspaced by a 100-ms blank screen. For each sentence, after presentation of the final segment, a test affirmation was shown (Table S1B) until the participant answered or for a maximum of 5 s. The participants were instructed to respond as quickly and correctly as possible via a button press using their left hand to indicate whether the affirmation was TRUE or FALSE with respect to the preceding sentence. The button-response association was counterbalanced across the participants.

One *syntax run* consisted of 16 trials featuring the two different syntactic structures (subject-relative and object-relative clauses) presented in a randomized order and in equal proportion (N = 8 trials each). The intertrial period was jittered between 4 and 6 s. The run started and finished with a 15 s baseline period. The sentences were visible through the mirror oriented towards the screen placed on the back of the scanner bore. The scripts controlling the presentation and recording of participants’ answers were programmed in *Psychtoolbox* (PTB-3, http://psychtoolbox.org/) running on MATLAB (Mathworks, Natick, USA) and built into a standalone executable thanks to the *deploytool* toolbox.

#### Motor Task

The participants performed a motor task inside the MRI scanner, requiring them to move a peg on a plastic board (Quercetti, Torino, Italy) between two fixed visual landmarks separated by an approximate 9-cm distance (Fig. 1D). The participants were required to perform this action with either their right hand (free-hand condition) or with a pair of 30-cm-long pliers held in their right hand (tool-use condition). One *motor run* consisted of 16 movements across the two conditions (tool-use and free-hand conditions) that were randomized (N=8 trials of each). One trial consisted in 5-s rest, another 5-s before the Go signal, followed by the movement phase (reach-to-grasp and object manipulation; Fig. 1D). The tool was initially placed on the chest of the participants, and they were prompted to catch it through the audio instruction “*pince”* (*pliers* in English). Instead, when receiving the audio instruction “*main”* (*hand* in English), the participants were prompted to place the pair of pliers back on their chest. The audio instruction to change the effector was only delivered when two consecutive trials required different effectors, otherwise without this audio instruction, the participants were prompted to keep using the same effector. When delivered, this audio instruction always occurred before the start of a new trial (i.e., before the 5-s rest period). After picking up the tool (tool-use condition) or returning the tool (free-hand condition) to their chest, the participants were prompted to place their hand (or tooled hand) at the starting position. Thus, in the tool-use condition, the participants already had the tool in their hand before starting a new trial. Once the hand (or tooled hand) was at the starting position, the trial started with the 5-s period followed by a single pure tone signal delivered through an MRI compatible device aiming to actively reduce MRI noise (Optoacoustics OptoACTIVE-two-way noise cancellation communication system, Mazor, Israel). After another 5 s, the movement phase was prompted by the double presentation of this same tone, indicating the Go for the action (see Fig. 1D). After having displaced the peg on the board, the participants were asked to return to the starting position so that a new trial could start. One motor run was preceded and terminated by a 15-s baseline period. If the participants left the starting position before the double tone (Go signal), an audio buzz was delivered prompting them to return to this position. These trials were considered as missed and were included in a regressor of non-interest for the fMRI analyses (1.30 % of trials). Furthermore, if a peg fell during the execution phase, the participants were prompted to grab a new peg from the left side of the plastic board. These events represented 0.55 % of the trials and the corresponding reach-to-grasp, object-manipulation and return phases were also considered as regressors of non-interest. The motor task device was placed in front of the participants at a reachable distance and made visible with a double mirror mounted onto the head coil. The participants’ right upper arm was strapped to the trunk to limit elbow and shoulder movements. To get reliable time measures of the motor behavior, three optic-fiber devices (FVDK 66, Baumer Electric AG, Frauenfeld, Switzerland) were mounted on the board to get information when the hand was on the starting position and when the peg was inserted within one of the two visual landmarks on the board. The output sent by the optic-fiber devices was collected through a parallel port controlled by the *io64* MATLAB function. The entire task was programmed in *Psychtoolbox* (PTB-3, http://psychtoolbox.org/) running on MATLAB (Mathworks, Natick, USA) and built into a standalone executable thanks to the *deploytool* toolbox.

### Procedure

The fMRI session was preceded by an inclusion session on a different day. This session aimed to describe the experiment and familiarize the participants with the tasks they underwent in the fMRI scanner. During this inclusion session, the participants performed a short version of the syntactic comprehension task (object-relative and subject-relative clauses, on different stimuli than the ones used for the fMRI session), and an adapted version of the motor task (tool and free hand). To ensure that enough trials could be analyzed after the fMRI session for the syntactic and motor tasks, cut-off performances were defined to include a participant. A total success rate of 62.5% (10 correct trials over 16) was expected for subject-relative clauses and 50% (8 correct trials over 16) for object-relative clauses. For the motor task, the participants were required to insert the first 10 pegs on a pegboard (Lafayette Grooved Pegboard Test), with the tool or the free hand (in two separate blocks for each effector). Participants were required on average to insert 10 pegs in less than 5-minutes with the tool and 1-minute with the free hand. Three participants that did not meet these criteria were excluded and could not take part in the fMRI session.

The fMRI session first consisted of a phase with syntactic and motor tasks inside the scanner (but without recordings) to familiarize the participants with the tasks in the MRI environment. This was followed by an anatomical (T1-weighted) and fieldmap acquisitions. Overall 14 runs were acquired in fMRI, including 7 for the motor task and 7 for the syntactic task. The runs were associated by pairs, with one motor run associated with a syntactic one, resulting in 7 pairs of runs. The order for each task within a pair was randomized within and between participants. A total of 224 trials were recorded over the whole session, consisting of 112 motor trials (with 56 trials for each tool-use and free-hand conditions) and 112 syntactic trials (with 56 trials for each object-relative and subject-relative conditions). Two participants did not complete the entire experiment, one did only 4 pairs of runs, while another one did only 5 of them. These subjects were still included in the analysis as the analyses were performed on session averages.

### Neuroimaging acquisition

Whole-brain structural and functional MRI scans were collected on a 3T (Siemens Prisma, Erlangen, Germany). The structural scan consisted of a T1-weighted (T1w) magnetization-prepared radio-frequency pulses and rapid gradient-echo (MPRAGE) scan (repetition time = 3000 ms, echo time = 3.8 ms, flip angle = 7°, matrix = 224 × 256, slice thickness = 1 mm, field-of-view = 224 × 256 mm, voxel size = 1 × 1 × 1 mm). Volumes were acquired with 192 interleaved slices. Functional scans instead consisted of a T2*-weighted (T2*w) gradient echo-planar imaging (EPI) sequence (3× multiband, repetition time = 1800 ms, echo time = 30 ms, flip angle = 80°, matrix = 112 × 112, slice thickness = 2 mm, 69 interleaved slices, field of view= 224 × 224 mm, voxel size = 2 × 2 × 2 mm).

### Analyses

#### Behavioral analyses

Linear mixed models (LMM) were used to compare the response times and sensitivity index (d′) between object-relative and subject-relative clauses (i.e., fixed-effect of conditions). The response times included only correct responses and were calculated from the test affirmation onset until the participants’ response. The subjects were included as random intercepts. For the response times, we also included the condition as random slope as the random matrix provided a better fit to the data[72]. We used the optimizer “bobyqa” to fit the LMMs, by estimating both fixed and random-effects parameters that maximized models’ likelihood. Random slopes could not be included in the model for the d′, as there was no within-subjects variability (i.e., only one observation per condition for each participant). The standard error of the mean (SEM) was calculated as a measure of dispersion between individuals.

#### fMRI preprocessing

The fMRI preprocessing was performed with fMRIprep, a tool that aims to offer reproducible ways to preprocess fMRI data[73]. The different steps applied during the preprocessing are precisely described in a file created by the fMRIprep pipeline and available online (https://osf.io/kyd72) In short, head motion parameters are estimated, then the images are corrected for magnetic field inhomogeneity from the fieldmap images. Following these steps, the images are slice-timing corrected, and co-registered to the T1w image in native space first and then in the MNI space. The fMRIprep pipeline estimated a number of time series, of which some are relevant to add to the subject-level models as nuisance regressors. For these models, we included the following time-series as nuisance regressors: the six motion parameters (i.e. translation and rotation across the Cartesian coordinates), the framewise displacement and the regressors estimated with the anatomical component-based noise correction method (aCompCor[74]). Regressors for the conditions of interest were also included in the model and are described in the *Subject-level analyses* section below. The same regressors of non-interest were used for both univariate and multivariate analyses (see below). fMRIprep does not apply any smoothing to the data.

#### Subject-level analyses

First-level analyses were performed with statistical parametric mapping (SPM12; Wellcome Trust Centre for Neuroimaging). Participant’s hemodynamic responses were modeled with a boxcar function. For the syntactic task, we modeled subject-relative and object-relative clauses during sentence presentation (i.e., syntactic encoding) and during test affirmation for correct trials separately. Incorrect trials were collapsed across the two syntactic conditions and were modelled for the sentence presentation and test affirmation phases separately. Sentence encoding was modeled for the entire presentation lasting 3.6 s, while the test affirmation was modeled for a duration corresponding to the participants’ RT for each single trial. For the motor task, we modeled tool use and free hand with three phases of interest: initiation, reach-to-grasp and object-manipulation phases separately, for correct trials only (see Fig. S1). The initiation phase was modeled from the 3-s period before participants reacted to the Go signal, ensuring to capture planning processes related to movement initiation (for similar procedure, see[75–77]). The reach-to-grasp phase started as soon as the participants started to move and ended when the peg was grabbed just before being pulled out from the board. The object-manipulation phase then ran from the time the peg was pulled out from the board until its insertion in a new position on the other side of the board. The onset and duration of each of these regressors depended on the participants’ performance in the task. Periods where the hand returned to the starting position or when effector change was instructed were modeled separately. We also modelled the 5-s rest period and the time preceding the initiation phase in distinct regressors. The rare incorrect trials when a peg fell or when the participants left the starting position too early were also modelled in the regressors of non-interest. All the regressors (of interest and non-interest) were the same for both univariate and multivariate analyses. The only difference was that for univariate analyses, the data was smoothed with a Gaussian kernel equal to three times the voxel size, resulting in a kernel with a full width at half maximum (FWHM) of 6mm^3^. For multivariate analyses, the data was not smoothed.

#### Group-level analyses

##### 1) Univariate analyses

At the second level, we conducted a paired sample t-test for the syntactic task, by contrasting the brain activity elicited by object-relative clauses with that elicited by subject-relative clauses in the encoding phase (β-values estimates across all runs). In the motor task, we contrasted the neural activity for tool use to that for free-hand actions in the three different phases: initiation, reach-to-grasp and object manipulation (β-values estimates across all runs). We reported as significant clusters passing cluster-level corrections for multiple comparisons set at P = 0.05 (cluster forming threshold set at P = 0.001 uncorrected). For the syntactic task, we further investigated the object-relative clauses activations with a region-of-interest (ROI) approach conducted on the left *pars opercularis* and the left *pars triangularis* of the inferior frontal gyrus (IFG). This analysis aimed to confirm that activity elicited by object-relative clauses (and subject-relative clauses) was different from baseline. The masks for these ROIs were obtained from the *freesurfer* brain parcellation implemented in *fmriprep*. Within each ROI, the β-values for object-relative and subject-relative clauses against baseline were extracted and averaged across voxels. A one-sided one-sample t-test was conducted to check whether the object-relative (and subject-relative) activations were different from zero in the left IFG. A further paired-sample t-test was run to corroborate the whole-brain analysis that activations for object-relative clauses did not differ from subject-relative clauses.

To assess the co-localization of neural resources between syntax and tool use, we employed a ROI approach. Based on the whole brain contrasts, we selected the syntactic ROIs involved in object-relative versus subject-relative clauses processing. Similarly, we defined the tool-use ROIs that were more strongly activated for tool-use than for free-hand actions, for each action phase separately. We thus selected the significant clusters derived from the univariate analyses presented in the previous paragraph. Only ROIs relevant for testing our hypotheses of co-localized neural resources between tool use and syntactic processes were considered. These included the left and right caudate nuclei clusters activated by object-relative clauses. No other clusters presented in Table 1 were deemed of interest with respect to our hypotheses (i.e., involvement of the basal ganglia). In the motor domain, we considered one cluster lying bilaterally within the caudate nuclei for tool-use reach-to-grasp (versus free-hand reach-to-grasp). We also considered two clusters in the left and right putamen activated by tool-use initiation (versus free-hand initiation). These two clusters were part of larger clusters extending to surrounding cortical areas but irrelevant to our hypotheses (see clusters 2 and 3 in Table 1B). Thus, we applied an anatomical mask (from *freesurfer*) to keep only the voxels in the basal ganglia (i.e., putamen). Within the ROIs extracted from the complex syntax contrast, we collected for each single voxel the β-values for each motor phase of tool-use and free-hand actions against the baseline. Instead, within the ROIs extracted from the tool-use contrasts, we collected the β-values for object-relative clauses and subject relative clauses versus baseline. Across voxels, averaged β-values were estimated for each ROI and participant, and they were submitted to a one-sided paired sample t-test, testing for higher neural activity for object-relative clauses (versus subject-relative clauses) or higher neural activity for tool use (versus free hand). A false rate discovery (fdr) rate was applied to correct for multiple comparisons and uncorrected p-values were also reported. Effect sizes were calculated with the Cohen’s d and SEM was calculated as a measure of dispersion between individuals.

##### 2) Multivariate analyses

Within the same ROIs (see Table 1 in bold), we tested how close/similar the neural patterns for object-relative clauses were to those for each tool-use phase (i.e., initiation, reach-to-grasp and object manipulation). We quantified the cosine similarity between the neural patterns for pairs of conditions. The cosine similarity lies in an interval between -1 and 1, where 1 means vectors are identical, while 0 means there is no relation between them. With a one-sided two-sample t-test, we tested whether the cosine similarity between the neural patterns for object-relative clauses and each tool-use phase was greater than the cosine similarity between the patterns for subject-relative clauses and each tool-use phase. We also tested whether the pattern similarity was higher between object-relative clauses and each tool-use phase, when compared to the similarity between object-relative clauses and each phase of the free-hand action. When appropriate, fdr corrections were applied to correct for multiple comparisons, in addition uncorrected p-values were reported. Effect sizes were calculated with the Cohen’s d and SEM was calculated as a measure of dispersion between individuals.

## Acknowledgments

We thank (in last name alphabetical order) Sophie-Anne Beauprez, Bertrand Beffara, Ivan Patané and Leslie Tricoche for their assistance during the MRI acquisitions. We are grateful to Danielle Ibarola and Franck Lamberton for their support in installing the experimental setup within the MRI facilities and for technical assistance during the MRI acquisitions. We thank Frédéric Volland for building some parts of the experimental device used.

## Notes

### Competing Interest Statement

The authors have declared no competing interest.

### Summary of Updates

Additions made to the discussion following Reviewer's feedbacks

