## Supporting Information for "Reaching for domain-general syntax: Sentence processing and tool-use reach-to-grasp share neural patterns in the basal ganglia"

### Supplementary Text

#### Methods - Behavior

To assess the existence of a learning effect, we analyzed the time course of the syntactic performance across the seven blocks. For both response times and d′, we used LMMs with the conditions (object-relative vs. subject-relative clauses) and the seven blocks as fixed effects. We employed a stepwise procedure to estimate which random effect to include in the model[1] . The main effect of conditions and blocks were included as random slopes and the subjects as random intercept. LMMs were also employed for the motor task to compare the performance between the tool-use and free-hand conditions in each of the three motor phases, namely for the initiation, reach-to-grasp and object-manipulation times. As for the syntactic task, we examined how performance evolved across the seven blocks for each of the three dependent variables. Both conditions (i.e., tool use vs. free hand) and blocks were used as fixed effects. Using the stepwise approach (as for syntax), we included the main effects of conditions and blocks as random slopes, while subjects were included as random intercepts.

#### Methods - fMRI

To corroborate the findings from the ROI-based approach, we performed a series of whole brain conjunction analyses (minimum statistic compared with conjunction-null hypothesis[2]) to assess the anatomical overlap between complex syntactic processing and tool use. Conjunction analyses are conservative because the same voxels need to pass the same threshold on two different maps to be considered as significant. Thus, we only reported clusters of more than 10 consecutives voxels at a threshold of P = 0.001 uncorrected (see[3]) .

#### Results - Behavior

In the MRI scanner, participants completed seven blocks of both the syntactic and motor tasks, progressively improving their performance over time. In the syntactic task, response times for correct answers significantly decreased from the first to the last block, regardless of sentence structure [block 1: 1888 ± 61 ms; block 7= 1626 ± 53 ms; $\chi^2_{(6)} = 24.08$, $P < 0.001$; Fig. S1A]. This learning effect was also reflected in the d′ sensitivity measure, which significantly increased from the first to the last block [block 1: 1.14 ± 0.07; block 7= 1.37 ± 0.05; $\chi^2_{(6)} = 13.21$, $P = 0.04$; Fig. S1B]. Similarly in the motor task, we observed participants improved from the first to the last block, irrespective of the effector (i.e. free hand or tool) employed for the action. The initiation time progressively decreased [block 1: 703 ± 10 ms; block 7= 638 ± 12 ms; $\chi^2_{(6)} = 21.71$, $P = 0.001$; Fig. S1C] as did the object-manipulation time [block 1: 1737 ± 66 ms; block 7= 1418 ± 50 ms; $\chi^2_{(6)} = 31.73$, $P < 0.001$; Fig. S1D]. However, for the reach-to-grasp time, we observed a different progression along the blocks depending on the effector [significant Effector x Block interaction: $\chi^2_{(6)} = 40.48$, $P < 0.001$; Fig. S1E]. At the beginning, the reach-to-grasp time difference between the tool and free-hand conditions was significantly greater [block 1 reach-to-grasp time difference = 523 ± 50 ms;

P < 0.001 fdr-corrected post-hoc tests] than at the end [block 7 reach-to-grasp time difference = 370 ± 51 ms; P < 0.001 fdr-corrected post-hoc tests], indicating greater initial difficulty in the tool condition that diminished with practice.

Results – fMRI

To confirm the overlap of neural activity observed between complex syntax processing and tool-use reach-to-grasp within the caudate nuclei (with the ROI-based approach), we ran a whole-brain conjunction analysis. This analysis tested for shared activations between syntax processing (object-relative > subject-relative clauses) and tool use (tool use > free hand) for the three action phases separately. For the reach-to-grasp phase of the tool-use action, we observed neural co-localization within the BG, specifically in the left posterior portion of the caudate nucleus head (uncorrected at P < 0.001; Fig. S4; Table S2). We did not find any overlap between syntax and tool use when considering the activations found during the tool-use initiation nor the object-manipulation phases.

**(A) Sentence Encoding**

| Subject-Relative Clauses (SRC) | Object-Relative Clauses (ORC) |
|---|---|
| L'écrivain qui admire le poète écrit le papier<br>The writer that admires the poet writes the paper<br>(Subject-object order compatible with the canonical order) | L'écrivain que le poète admire écrit le papier<br>The writer that the poet admires writes the paper<br>(Noncanonical subject-object order) |

**(B) Test Affirmation (one selected among the four)**

| | |
|---|---|
| L'écrivain admire le poète<br>The writer admires the poet<br>(SRC = True - ORC = False) | Le poète admire l'écrivain<br>The poet admires the writer<br>(SRC = False - ORC = True) |
| Le poète écrit le papier<br>The poet writes the paper<br>(SRC = False - ORC = False) | L'écrivain écrit le papier<br>The writer writes the paper<br>(SRC = True - ORC = True) |

**Table S1**. Syntactic task: comprehension of sentences in a 2-alternative forced choice (2-AFC) task. (A) Syntactic structures presented during the sentence-encoding phase. (B) Test affirmation for the 2-AFC task (2 true and 2 false possible probes for each encoded sentence).

| Region | BA | Peak MNI coordinates | | | Clust er size | Z-value |
|---|---|---|---|---|---|---|
| | | x | y | z | | |
| **Conjunction: Syntactic network (A) ∩ Tool-use network – reach-to-grasp (C)** | | | | | | |
| L Caudate Nucleus | - | -6 | 8 | 0 | 18 | 4.13 |

**Table S2**. Conjunction between the syntactic network and the tool-use network for the reach-to-grasp phase. From left to right are reported the names of the regions in which the clusters were identified, the Brodmann Area (BA), the Montreal Neurological Institute (MNI) coordinates for the peak of activity, the cluster size in number of voxels and the statistics with the Z-values. The clusters highlighted in bold were used for regions-of-interest (ROIs) analyses.

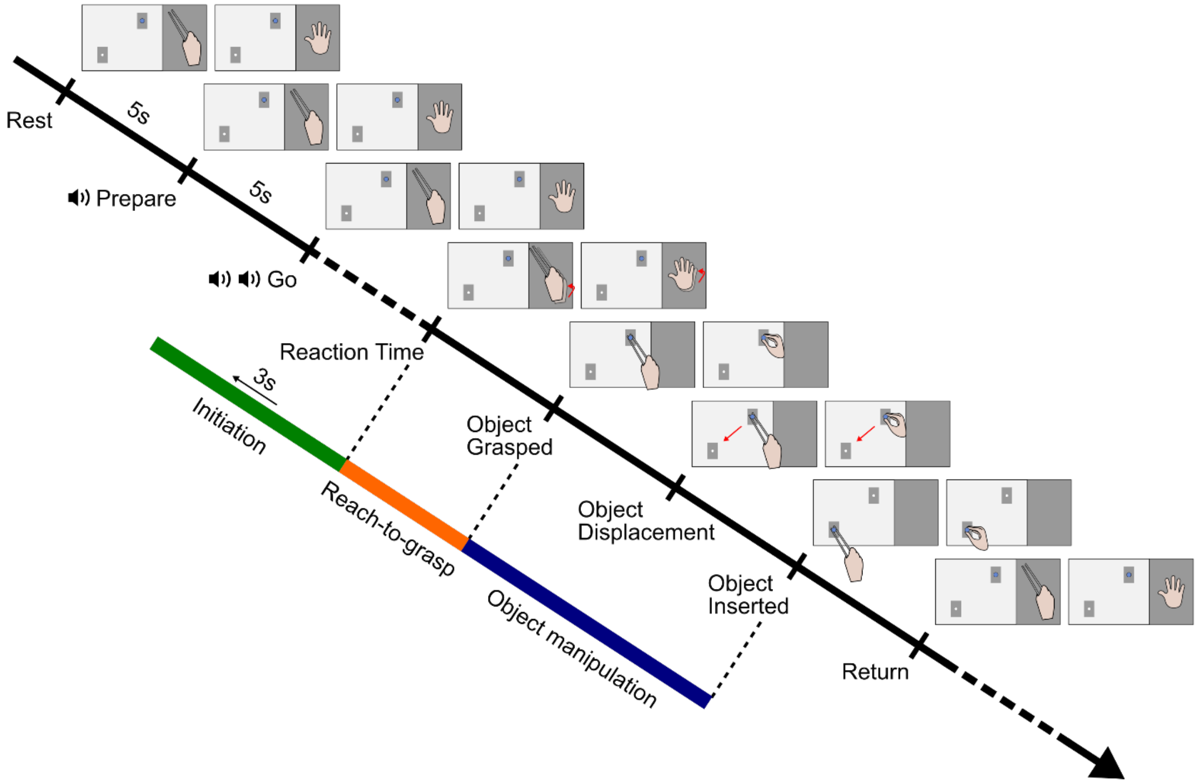


**Figure S1.** Motor sequence executed with the free hand (top row) and tool (bottom row). Participants started with a “Rest” period followed by the “Preparation” period. The “Go” signal prompted the participants to reach and grasp the target object (i.e., blue peg) before the manipulation (i.e., moving the target object from its initial to final position). Participants were finally asked to return to the start position with their hand (i.e., dark grey platform) to prepare for the next trial. The initiation (green), reach-to-grasp (orange) and object-manipulation phases (blue) were modelled as regressors of interest for the fMRI analyses (see Methods). The initiation phase included the 3-second period preceding the onset of the movement (i.e. time-locked to the participants’ reaction time). Next, the reach-to-grasp phase included the time from the first motion of the hand to the grasping of the target object. Finally, the object-manipulation phase ran from the target-object grasping to its insertion in the final position.

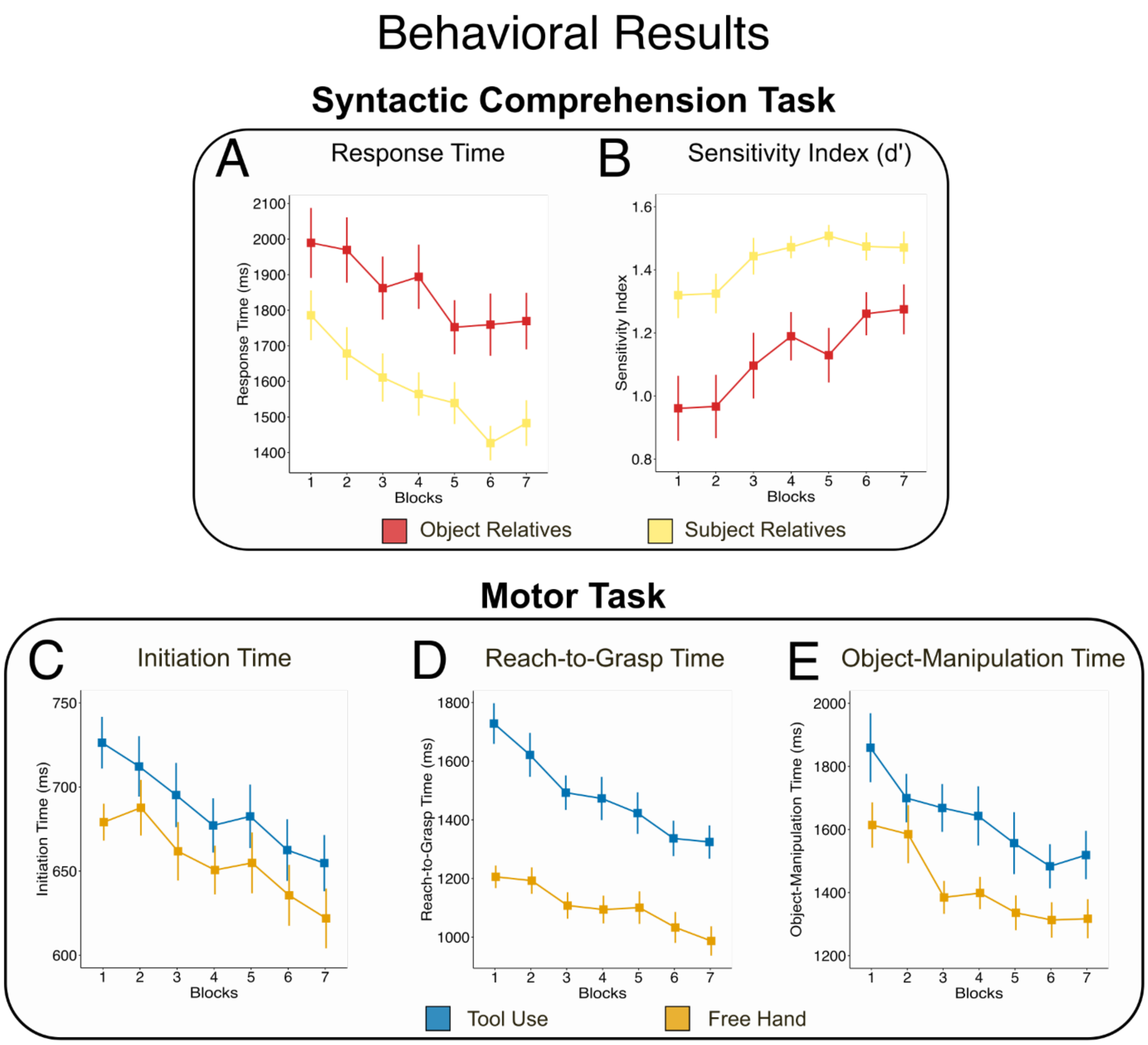


**Figure S2.** Syntactic performance across blocks: response times (A) and sensitivity index (B). Object-relative clauses are in red and subject-relative clauses in yellow. Both the response time and sensitivity index show a main effect of condition, with slower response times and reduced sensitivity index for the comprehension of object-relative clauses. Motor performance across blocks: initiation time (C), reach-to-grasp time (D), object-manipulation time (E). Tool use is in blue and free hand in orange. For each phase, there is a main effect of condition, with longer times for tool use as compared to free hand. For the reach-to-grasp phase, there is also an interaction between the conditions and the blocks, given by the reduction across blocks of the difference between the two conditions. The error bars represent the standard error of the mean (SEM).

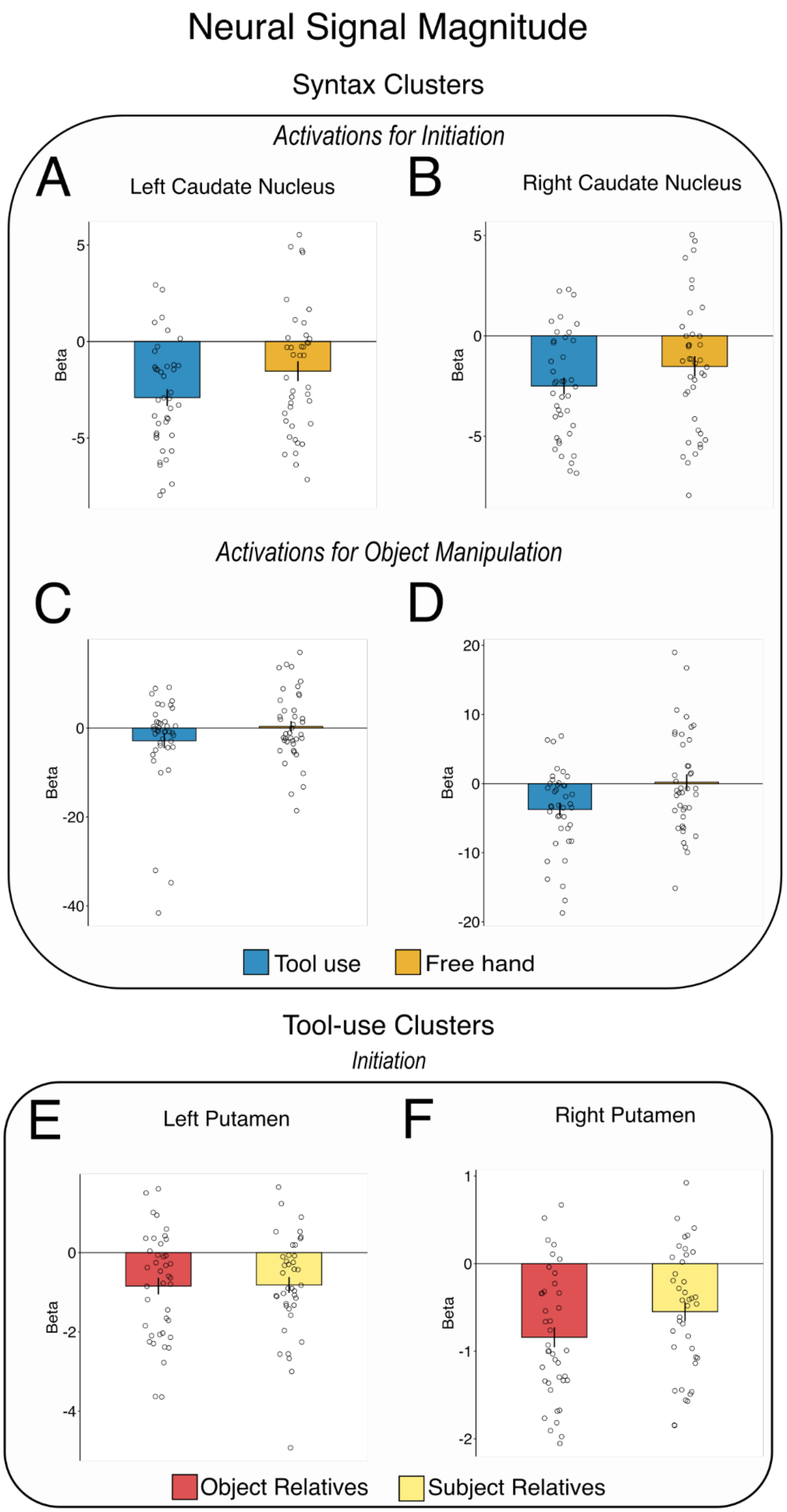
Neural Signal Magnitude
Syntax Clusters
Activations for Initiation
A
Left Caudate Nucleus
B
Right Caudate Nucleus
Activations for Object Manipulation
C
D
Beta
Tool use
Free hand
Tool-use Clusters
Initiation
E
Left Putamen
F
Right Putamen
Object Relatives
Subject Relatives

**Figure S3.** Neural activations for the motor initiation phase in the clusters activated by complex syntax (object-relative vs. subject-relative clauses): left caudate nucleus (A) and right caudate nucleus (B). Neural activation for the object-manipulation phase in the same syntactic clusters: left caudate nucleus (C) and right caudate nucleus (D). Tool use is in blue and free hand in orange. Neural activations for syntax in the clusters activated by tool-use initiation: left putamen (E) and right putamen (F). Object-relative clauses are in red and subject-relative clauses in yellow. The error bars represent the standard error of the mean (SEM).

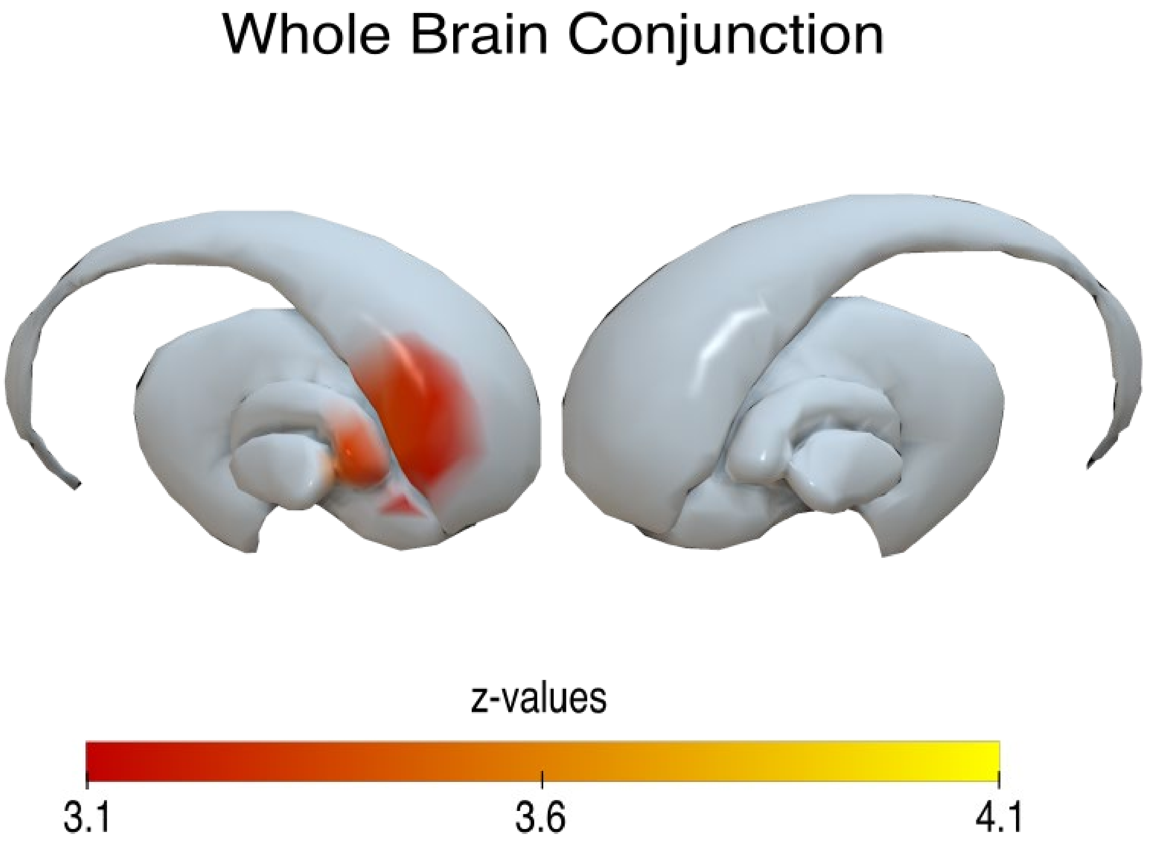


**Figure S4.** Through a whole brain analysis, we found an overlap between syntax and tool-use reach-to-grasp in the basal ganglia, mainly in the left caudate nucleus (D). This map was thresholded with a voxel-wise threshold at P = 0.001. The localization of the activations is described in Table S2.

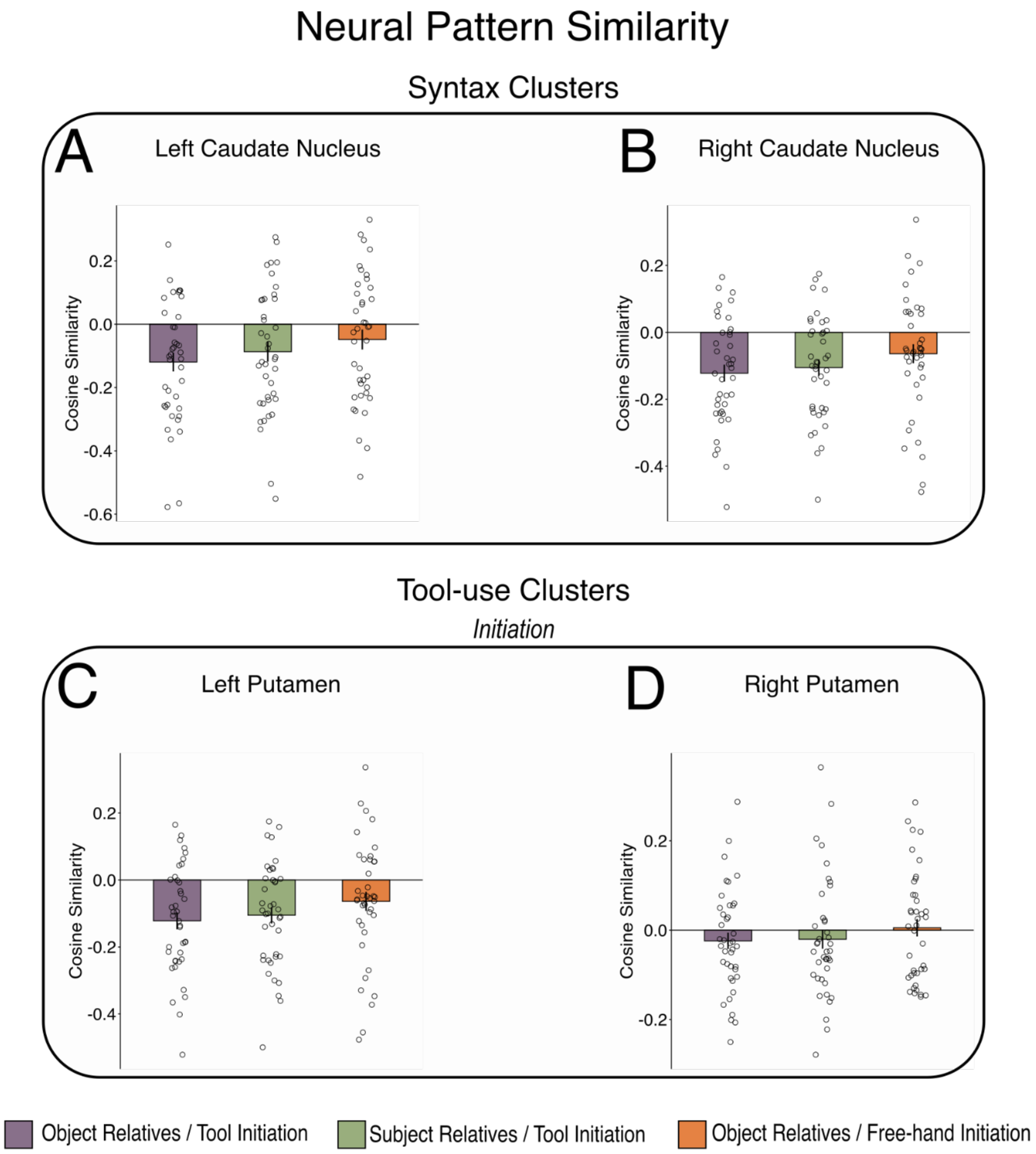


**Figure S5.** Neural pattern similarities between syntax and motor activations for the initiation phase, in the left caudate nucleus (A) and the right caudate nucleus (B) activated by complex syntax (object-relative vs. subject-relative clauses), and in the left putamen (C) and right putamen (D) activated by tool-use initiation (vs. free-hand initiation). Neural pattern similarity is measured with the cosine distance for the three pairs: object-relative clauses and tool-use initiation in purple, subject-relative clauses and tool-use initiation in green and object-relative clauses and free-hand initiation in orange. The error bars represent the standard error of the mean (SEM).

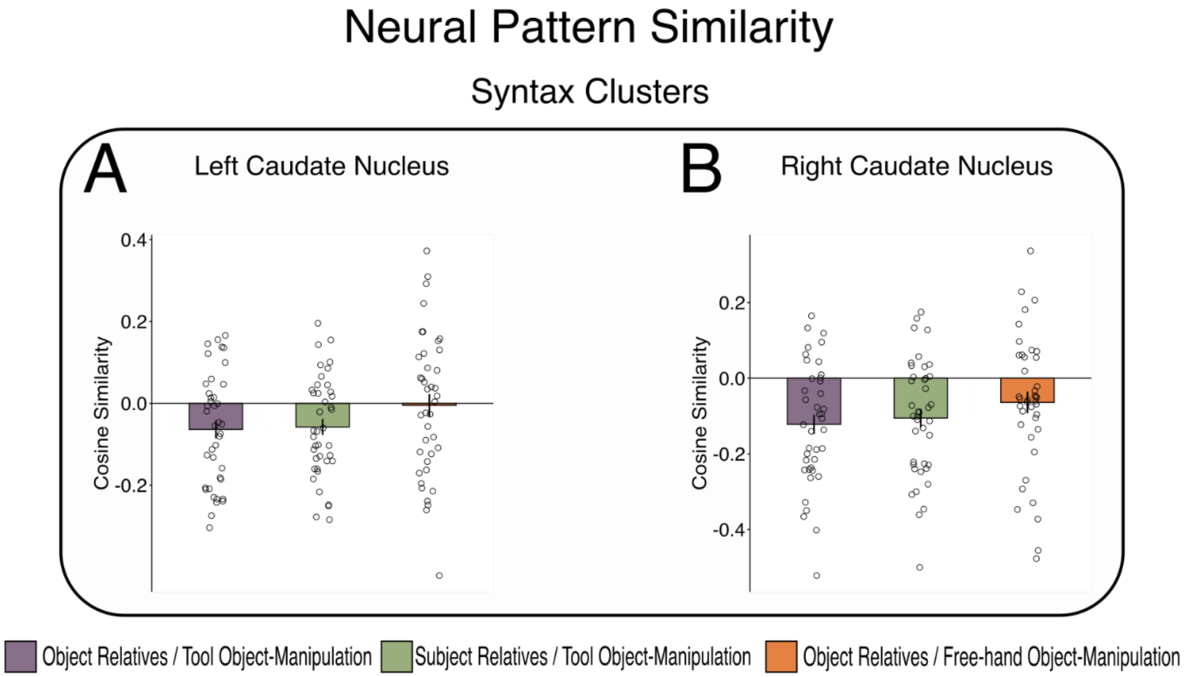


**Figure S6.** Neural pattern similarities between syntax and motor activations for the object-manipulation phase, in the left caudate nucleus (A) and the right caudate nucleus (B) activated by complex syntax. Neural pattern similarity is measured with the cosine distance for the three pairs: object-relative clauses and tool-use object-manipulation in purple, subject-relative clauses and tool-use object-manipulation in green and object-relative clauses and free-hand object-manipulation in orange. The error bars represent the standard error of the mean (SEM).

**Video S1.** Motor task in fMRI scanner. This video depicts object manipulation with and without a tool (i.e., pair of pliers).
